# OTUB1 controls marginal zone B-cell development by stabilizing RelA in a CD40-dependent manner

**DOI:** 10.64898/2026.08.06.743158

**Authors:** Johannes F. Vogt, Yilang Tang, Sonja Reißig, Christina Karantanou, Shashank Kumar, Emmanouil Stylianakis, Dirk Schlüter, Ari Waisman, Nadine Hövelmeyer

## Abstract

Ubiquitin-dependent regulation of NF-κB signaling is essential for B-cell homeostasis and fate decisions, yet the contribution of specific deubiquitinating enzymes remains incompletely defined. OTUB1, a lysine-48–specific deubiquitinase, has been broadly implicated in immune regulation, including control of NF-κB signaling and prevention of immune hyperactivation. Previous studies have demonstrated that B cell-specific deletion of OTUB1 leads to B cell hyperplasia, increased antibody production, and lupus-like autoimmunity, highlighting its importance in maintaining B cell tolerance and immune homeostasis. Using B cell-specific OTUB1-deficient mice, we show that loss of OTUB1 leads to a marked expansion of marginal zone (MZ) B cells and their precursor populations in the spleen, accompanied by an activated phenotype and enhanced proliferative responses, particularly upon CD40 stimulation. OTUB1 deficiency results in altered CD40-induced NF-κB signaling, characterized by enhanced IκB degradation and increased nuclear accumulation of p50-containing NF-κB complexes, despite reduced RelA stability. Mechanistically, we show that OTUB1 interacts with RelA, restricting its lysine-48–linked ubiquitination and proteasomal degradation, thereby stabilizing this key transcription factor. Collectively, these findings identify RelA as a novel OTUB1 target and uncover an additional layer of ubiquitin-dependent control of NF-κB signaling that governs splenic B-cell homeostasis and marginal zone B-cell development.

## Introduction

Naïve splenic B cells consist of immature-transitional (T) B cells and three types of mature B cells: follicular (FO), marginal zone (MZ) B cells, and B1 cells. Immature B cells that emigrate from the bone marrow and enter the spleen develop through distinct transitional B cell stages, termed transitional (T)1, T2, and T3 (1). The T2 B cells are thought to be the common precursors for both MZ and FO B cells, which differ in their surface phenotype, anatomic localization, and immunologic function(Allman & Pillai, 2008). A variety of signals have been shown to influence the cell fate decision between FO and MZ B cells, including signals provided by the family of Notch receptors (Gibb et al., 2010; Maillard et al., 2004), the B cell receptor (BCR) (Casola et al., 2004; Heltemes & Manser, 2002), and NF-κB signaling pathways (Franzoso et al., 1997; Grossmann, 2000).

The NF-κB family of transcription factors consists of structurally related proteins, namely RelA (p65), c-Rel, RelB, p105/p50 (NF-κB1), and p100/p52 (NF-κB2) (Hövelmeyer et al., 2022). In mature B cells, NF-κB-activating signals can be delivered from immune cell-surface receptors such as the BCR, Toll-like receptors (TLRs), and TNF family receptor (TNFR), including CD40 and B-cell activating receptor (BAFFR). These receptors transduce cellular signals to govern the physiological and pathological processes in B cells, including B cell development and differentiation, survival, proliferation, and antibody-mediated immune responses, as well as autoimmune diseases and B cell lymphomagenesis (Chen & Wang, 2021). To prevent persistent activation of NF-κB, a tight control of this pathway is required. This is achieved, amongst other mechanisms, by post-translational modifications, such as phosphorylation and ubiquitination.

Protein ubiquitination is a process involving the covalent fusion of one or more ubiquitin monomers that can be linked via different lysine residues. Depending on their type of ubiquitination, proteins may be degraded by the proteasome (lysine 48) or functionally altered during signal transduction (lysine 63). Ubiquitination is a reversible process, and the degree of ubiquitination of specific proteins is controlled by the concerted actions of conjugating enzymes and deubiquitinating enzymes (DUBs). Several deubiquitinating enzymes have been reported, by us and others, to negatively regulate NF-κB signaling, particularly in B cells, amongst them A20 (Chu et al., 2011; Hövelmeyer et al., 2011; Tavares et al., 2010), CYLD (HoÖvelmeyer et al., 2007), Cezanne (Luong et al., 2013), and OTUB1 (Li et al., 2019).

OTUB1 (OUT domain ubiquitin aldehyde-binding protein 1) belongs to the ovarian tumor domain proteases (OUT) subfamily of DUBs and negatively regulates ubiquitination to control protein stability and activity (Edelmann et al., 2009; He et al., 2017; Komander et al., 2009). OTUB1 preferentially cleaves lysine-48-linked poly-ubiquitin chains and prevents ubiquitination by blocking ubiquitin transfer to target proteins, such as Snail(Zhou et al., 2018), Smad2/3 (Herhaus et al., 2013), UBC13 (Mulas et al., 2021), and p100 (Li et al., 2019), and thereby plays an important regulatory role at the level of protein turnover by preventing degradation of these target proteins. Cell-type-specific deletion of *Otub1* in immune cells and other tissues confirmed its critical role in the regulation of different signaling pathways, including GPCR and NF-κB signaling. Li *et al*. identified OTUB1 as a p-100-interacting protein that inhibits the ubiquitination and degradation of p100. They showed that OTUB1 deficiency caused spontaneous ubiquitination and degradation of p100 in mouse embryonic fibroblasts and B cells. Moreover, OTUB1 deficiency prevents accumulation of p100 with canonical NF-κB activation, leading to aberrant activation of both canonical and noncanonical NF-κB members (Y. Li et al., 2019). In a more recent study, Luo *et al*. (Luo et al., 2024) discovered a novel role for OTUB1 as a regulator of G protein-coupled receptor signaling in B cells, with its deficiency affecting B cell differentiation, including both marginal zone and follicular B cells, as well as their splenic localization.

In the present work, we unraveled an additional, so far unknown role of OTUB1 in B cell homeostasis and NF-κB activation. We show that the conditional deletion of OTUB1 in B cells results in a dramatic increase in MZ B cells. These B cells display increased class switch recombination capacity and have an increased proliferative ability after αCD40 stimulation.

Mechanistically, we show that OTUB1 regulates the kinetics of CD40-induced NF-κB signaling by stabilizing RelA. Loss of OTUB1 promotes K48-linked ubiquitination and proteasomal degradation of RelA and is accompanied by increased nuclear p50 and altered IκB degradation after αCD40 stimulation

## Results and Discussion

### OTUB1 Limits Marginal Zone B Cell Differentiation

To investigate the B cell-specific role of OTUB1, OTUB1^FF^ mice were crossed to *Cd19*-cre mice (thereafter named: OTUB1^BKO^). Macroscopic examination of secondary lymphoid organs revealed no differences in the size of the spleens compared to *Cd19*-cre control animals (**Fig. 1A**). Histological analysis of the spleen further revealed no obvious differences in splenic architecture between *Cd19*-cre control and OTUB1^BKO^ mice (**Supplementary Fig. 1A**). Total splenic cell numbers (**Fig. 1B**) as well as B and T-cell numbers were comparable to age-matched controls (**Supplementary Fig. 1B**).

**Figure 1:**
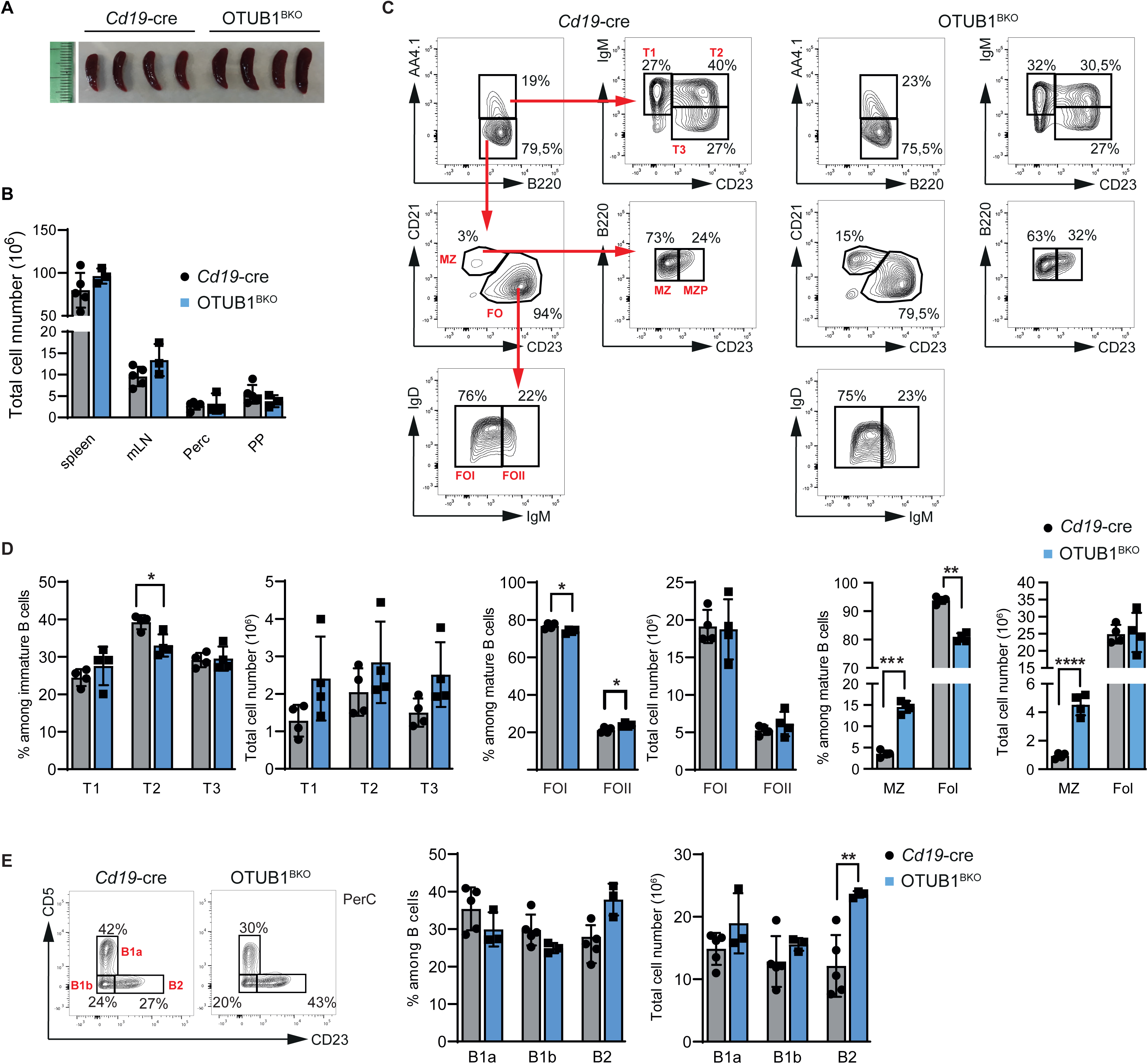
OTUB1 regulates B cell differentiation and subset distribution. **(A)** Representative images of spleens from control (Cd19-cre) and B cell–specific Otub1-deficient (Otub1BKO) mice. **(B)** Total cell numbers in spleen, mesenteric lymph nodes (mLN), peritoneal cavity (PerC), and Peyer’s patches (PP) from Cd19-cre and Otub1BKO mice. **(C)** Representative flow cytometry plots showing gating strategy and frequencies of splenic B-cell subsets. Transitional B cells (T1, T2, T3) were identified based on AA4.1, B220, CD23, and IgM expression. Mature B-cell subsets, including follicular (FO), marginal zone (MZ), and marginal zone precursor (MZP) B cells, were defined using CD21 and CD23. **(D)** Quantification of transitional (T1–T3), follicular (FOI, FOII), and marginal zone (MZ) B-cell subsets in Cd19-cre and Otub1BKO mice, shown as frequency and total cell number. **(E)** Representative flow cytometry plots and quantification of peritoneal cavity B-cell subsets, including B1a (CD5⁺), B1b (CD5⁻), and B2 cells, in Cd19-cre and Otub1BKO mice. Data are shown as mean ± SEM. Each dot represents an individual mouse. n = 3 - 5 mice of each genotype, and are representative of at least three independent experiments. Statistical significance was determined using unpaired two-tailed Student’s t-test, with *p < 0.05, **p < 0.01, ***p < 0.001, ****p < 0.0001.

To enumerate the different B cell subpopulations in 8–12-week-old OTUB1^BKO^ mice, we performed flow cytometric analysis. Cell surface expression of CD21 and CD23 discriminates between immature (CD21^-^CD23^-^), FO (CD21^int^CD23^high^), and MZ (CD21^high^CD23^low^) B cells. The percentage of splenic immature (AA4.1^+^) B cells was not significantly changed (**Fig. 1C**). Further analyses of immature splenic B cells also showed no major changes in transitional 1 (T1) (IgM^+^CD21^low^CD23^low^) and T3 (IgM^+^CD21^high^CD23^high^) cells whereas the frequency of T2 (IgM^+^CD21^high^CD23^high^) cells among immature B cells was modestly increased in OTUB1^BKO^ mice, however, absolute numbers of T1-T3 were not significantly different form the *CD19-Cre* controls (**Fig. 1C and D**). Among mature B cells, OTUB1^BKO^ mice showed significant shifts within the follicular compartment, with FOI (IgM^int^, CD23^high^) cells being reduced and FOII (IgM^high^, CD23^high^) cells relatively enriched in frequency, while total FOI and FOII cell numbers remained unchanged (**Fig.1C and D**). In contrast, OTUB1^BKO^ mice displayed a profound and significant increase in the relative and absolute numbers of MZ B cells (B220^+^AA4.1^-^CD21^hi^CD23^low^) while total cell counts of FO B cells were not significantly changed compared to *Cd19*-cre controls (**Fig.1C and D**). A similar expansion of MZ B cells accompanied by reduced FO representation was reported in Mb1-Cre OTUB1^FF^ mice, although in that model FO B cells were decreased in both frequency and absolute number (Luo et al., 2024), whereas in our Cd19-Cre line FO cell numbers remain preserved at 8–12 weeks. Thus, loss of OTUB1 in CD19-expressing B cells specifically leads to a selective alteration to the splenic MZ B cell subset, while remodeling the composition of the follicular department without causing a global loss of FO B cells. Finally, in an additional cohort of 30-week-old OTUB1^BKO^ mice, we observed that the skewing in subset frequencies persisted and was even more pronounced, with further increased MZ and decreased FO proportions among splenic B cells **(Supplementary Fig. 1C)**.

Within the peritoneal cavity (PerC), B cells are classified into B1 and B2 subsets(Hardy & Hayakawa, 2015), with B1 cells presenting the predominant population. B1 cells can be further subdivided into B1a and B1b subsets, distinguished by the expression of CD5 on B1a cells, whereas B1b cells lack CD5 expression (Duan & Morel, 2006). Flow cytometric analysis of the PerC revealed that OTUB1 deficiency increased the frequency of B2 cells, while the frequency as well as total numbers of B1a and B1b cells were not significantly changed compared to controls (**Fig. 1E**). Thus, OTUB1 deficiency results in a specific increase of the MZ B cell population, but no other B cell type.

### OTUB1 modulates anti-CD40-driven B cell activation

To analyze the role of OTUB1 in the proliferative capacity of B cells as a response to individual and combined stimuli, B cells were labeled with VCT, cultured with or without LPS, αIgM, CpG, αCD40, or αCD40+IL-4 followed by flow cytometric analysis after 3 days (**Fig. 2A**). While αIgM, LPS, and CpG-induced proliferation was comparable to control B cells, OTUB1-deficient B cells stimulated with αCD40 alone or with αCD40+IL-4 showed increased proliferation compared to control B cells (**Fig. 2A**). Thus, OTUB1 seems to be specifically involved in αCD40 driven B cell activation.

**Figure 2:**
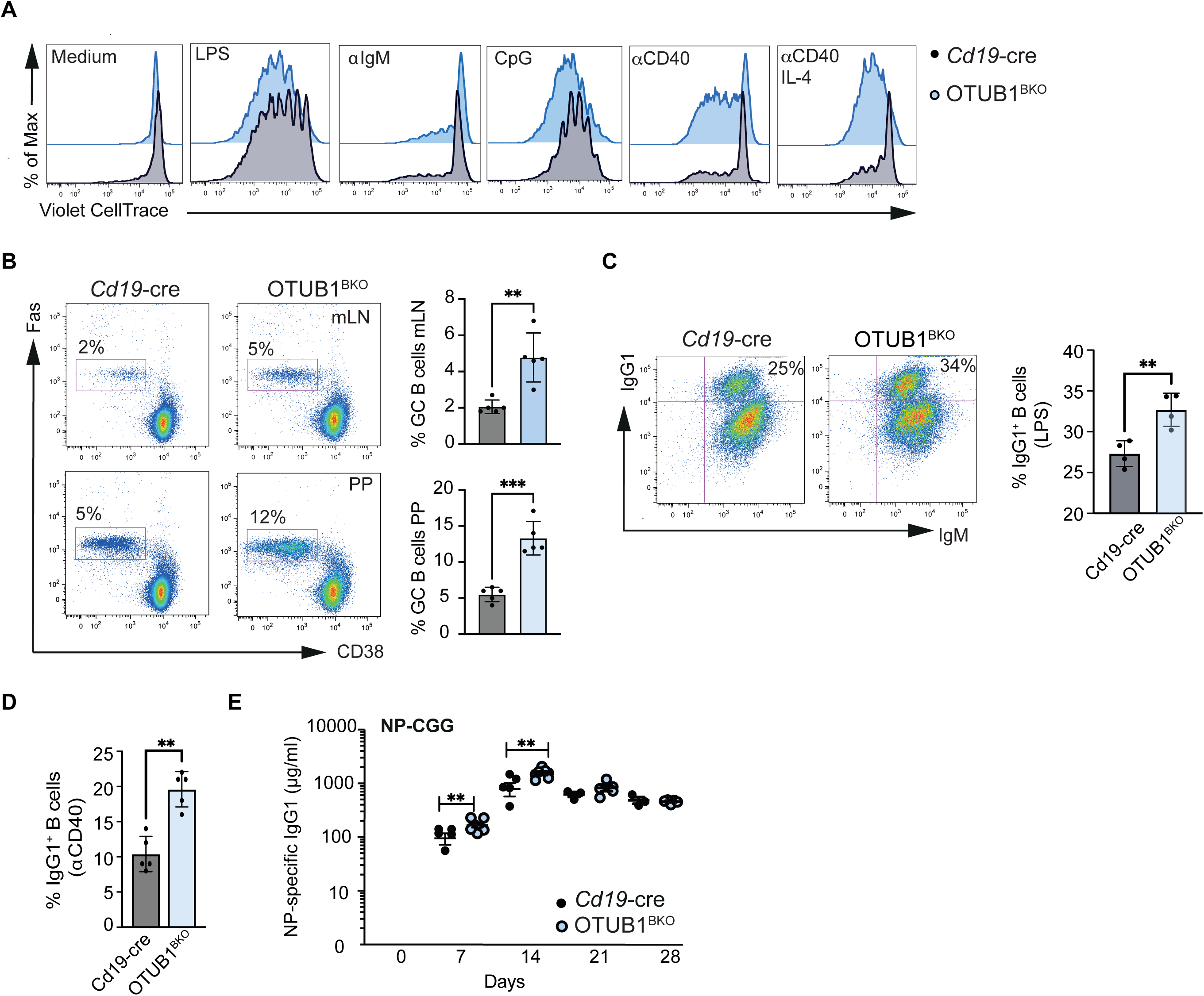
OTUB1 restrains T-cell-dependent B-cell activation and class-switch responses. **(A)** Proliferation of MACS-purified LN B cells from control (Cd19-cre) and B cell–specific Otub1-deficient (Otub1BKO) mice assessed by CellTrace Violet (VCT) dilution following stimulation with medium, LPS, αIgM, CpG, αCD40, or αCD40 plus IL-4 for 72 hours. Representative histograms are shown. **(B)** Representative flow cytometry plots and quantification of germinal center (GC) B cells (Fas^+^CD38^-^) in mesenteric lymph nodes (mLN) and Peyer’s patches (PP) from Cd19-cre and Otub1BKO mice. Percentages of GC B cells are indicated. **(C)** Representative flow cytometry plots and quantification of IgG1 class switching in MACS-purified LN B cells stimulated with LPS for 4 days. Percentages of IgG1⁺ B cells are indicated. **(C)** Quantification of IgG1⁺ B cells following in vitro stimulation with αCD40 for 4 days. CD43-depleted lymph node B cells were analyzed by flow cytometry for class switch recombination. **(D)** NP-specific IgG1 serum antibody titers measured by ELISA following immunization with NP-CGG in Cd19-cre and Otub1BKO mice over time (days 7, 14, 21, and 28). Data are shown as mean ± SEM. Each dot represents an individual mouse. All experiments were performed in at least three independent experiments. Statistical significance was determined using unpaired two-tailed Student’s t-test (panels A–D) or multiple unpaired t-tests with Welch’s correction (panel E), with *p < 0.05, **p < 0.01, ***p < 0.001, ****p < 0.0001.

CD40 signaling of B cells promotes the formation of long-lived plasma cells and memory B cells, as well as germinal center (GC) formation and immunoglobulin (Ig) switching (Danese et al., 2004). Indeed, we found a significant increase in the percentage of GC B cells in the mesenteric lymph nodes (mLN) and Peyer’s patches (PP) compared to control animals (**Fig. 2B**). Next, we isolated B cells and analyzed them for their class switch to IgG1-expressing cells. As shown in **Fig. 2C** and **D**, OTUB-deficient B cells exhibited a significantly greater capacity to class-switch to IgG1^+^ B cells under both conditions than B cells from *Cd19*-cre mice. To determine how OTUB1^BKO^ mice respond to T-cell-dependent antigens *in vivo*, we immunized OTUB1^BKO^ and control mice with NP-CGG (4-hydroxy-3-nitrophenylacetyl–chicken gamma globulin) to induce a B-cell response. Measurement of NP-specific antibody titers at different time points after immunization showed significantly increased amounts of NP-specific IgG1 in the sera of immunized OTUB1^BKO^ mice at 7 and 14 days after immunization, with a peak at day 14, and then remained elevated and comparable between the two groups at later time points (**Fig. 2E).** By contrast, naive serum immunoglobulin concentrations were comparable between *Cd19*-cre and OTUB1^BKO^ mice under steady-state conditions **(Supplementary Fig. 1D),** indicating that the enhanced humoral phenotype becomes most apparent upon stimulation. Taken together, these findings indicate that OTUB1 negatively regulates CD40-dependent proliferation as well as class-switch responses in B cells.

### OTUB1 limits K48-linked ubiquitination and degradation of RelA

Because CD40-induced proliferation critically depends on the activation of NF-κB (Berberich et al., 1994; Coope, 2002), and we observed increased proliferation upon αCD40, the canonical and non-canonical pathways were analyzed by Western blot. Cytoplasmic and nuclear fractions were isolated from LN B cells, which were either left untreated or stimulated with αCD40 for the indicated time points.

In contrast to the findings reported by Li et al., we did not observe obvious differences in p100 protein abundance between *Cd19*-cre and OTUB1^BKO^ B cells at baseline or during *⍺*CD40 stimulation (**Fig. 3A**). Likewise, p100 phosphorylation and processing to p52 were not altered (**Fig. 3A**), suggesting that OTUB1-dependent regulation of p100 may be stimulus- or context-dependent. Consistent with this, NF-κB signaling components, including p100, p52, RelA, and RelB, were not altered in OTUB1^BKO^ cells stimulated with BAFF (**Supplementary Fig. 2A)** or LPS **(Supplementary Fig. 2B).** Interestingly, RelA protein levels were reduced in OTUB1^BKO^ B cells after prolonged αCD40 stimulation. Specifically, we observed a marked loss of RelA in the cytoplasmic fraction and an almost complete absence of nuclear RelA accumulation at 8 and 16 h after αCD40 stimulation (**Fig. 3A**). Although p105 protein abundance was not altered in OTUB1^BKO^ B cells after αCD40 stimulation, we detected increased nuclear p50 after 8 and 16h after αCD40 stimulation (**Fig. 3B**).

**Figure 3:**
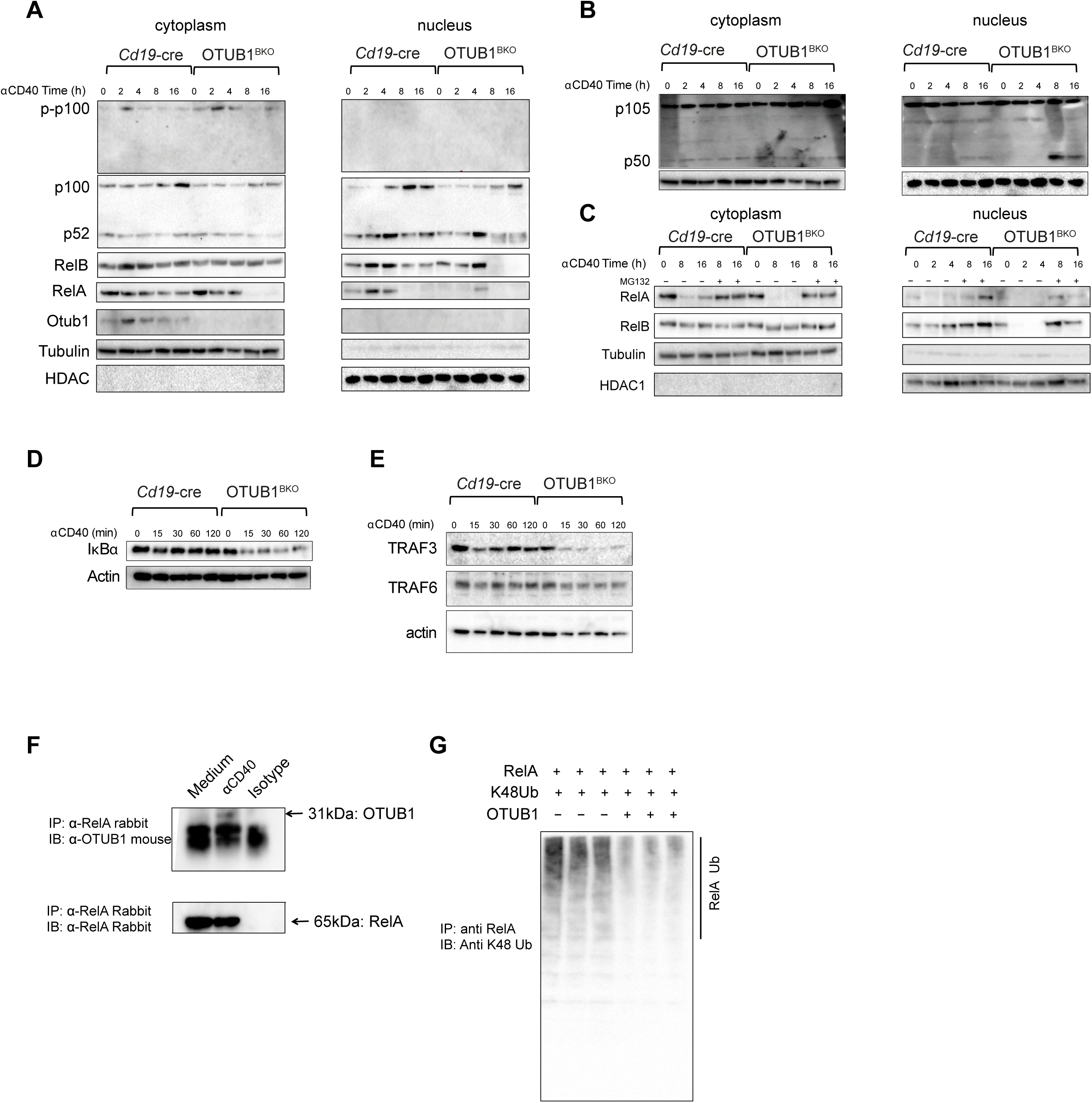
OTUB1 stabilizes RelA after CD40 stimulation. **(A)** Immunoblot analysis of cytoplasmic and nuclear fractions from B cells isolated from Cd19-cre and Otub1BKO mice stimulated with αCD40 for the indicated time points (0, 2, 4, 8, 16 h).. Blots show levels of phosphor-p100 (p-p100), p100, p52, RelA, and RelB. Tubulin and HDAC1 serve as cytoplasmic and nuclear loading controls, respectively. **(B)** Immunoblot analysis of cytoplasmic and nuclear extracts showing p105 and p50 levels following αCD40 stimulation (0, 2, 4, 8, 16 h) in Cd19-cre and Otub1BKO B cells.**(C)** Immunoblot analysis of RelA and RelB expression in cytoplasmic and nuclear fractions following αCD40 stimulation (8 and 16 h) in the presence (+) or absence (−) of the proteasome inhibitor MG132 (10 μM) Tubulin and HDAC1 serve as loading controls. **(D)** Immunoblot analysis of IκBα degradation in B cells stimulated with αCD40 for the indicated time points (0, 15, 30, 60, 120 min). Actin serves as a loading control. **(E)** Immunoblot analysis of TRAF3 and TRAF6 expression following αCD40 stimulation for the indicated time points (0, 15, 30, 60, 120 min). Actin serves as a loading control. **(F)** Co-immunoprecipitation analysis showing interaction between RelA and OTUB1 in B cells under basal conditions and following αCD40 stimulation. Cell lysates were immunoprecipitated (IP) with anti-RelA antibody or control rabbit IgG, followed by immunoblotting (IB) with anti-OTUB1 or anti-RelA antibodies. **(G)** In vitro ubiquitination assay showing K48-linked ubiquitination of RelA in the presence or absence of OTUB1. RelA was immunoprecipitated and probed with anti–K48-linked ubiquitin antibodies. Data are representative of at least 2-3 independent experiments.

To test whether the loss of RelA in OTUB1KO B cells was due to enhanced proteasomal degradation, we stimulated B cells for 8 and 16 h with αCD40 in the presence or absence of the proteasomal inhibitor MG132. MG132 treatment restored, at least in part, RelA protein levels after αCD40 stimulation in OTUB1^BKO^ cells, supporting the idea that OTUB1 limits proteasome-dependent RelA degradation. (**Fig. 3C**). Although the most prominent difference observed was the loss of RelA after prolonged CD40 stimulation, additional changes in upstream signaling components were also apparent. Specifically, OTUB1-deficient B cells showed altered I*Κ*B*⍺* kinetics after CD40 stimulation, as evidenced by differences in phosphorylated and total I*Κ*B*⍺* levels (**Supplementary Fig. 2C and Fig. 3D**). By contrast, *⍺*IgM-induced I*Κ*B*⍺* degradation was broadly comparable between Cd19-cre and Otub1KO B cells (**Supplementary Fig. 2D**). In addition, because TRAF2, TRAF3, and TRAF6 are important mediators of CD40 signal transduction (Bishop et al., n.d.), and OTUB1 and OTUB2 were shown to regulate TRAF3 and 6 stabilities (S. Li et al., 2010), we aimed to investigate these two proteins after CD40 activation. Interestingly, we did detect a decrease in the protein level of TRAF3 after aCD40 stimulation of the OTUB1-deficient B cells, while TRAF6 protein levels were almost unchanged (**Fig. 3E**). Finally, to further examine whether OTUB1 directly binds to RelA we performed an immunoprecipitation experiment in B cells under basal conditions or after *⍺*CD40 stimulation. As can be seen in **Fig. 3F**, OTUB1 indeed binds to RelA after αCD40 stimulation, identifying RelA as a previously unappreciated target of OTUB1. Together with the MG132 rescue experiments, these findings suggest that OTUB1 protects RelA from proteasome-dependent degradation during CD40-induced signaling.

Since OTUB1 is a deubiquitinating enzyme, we tested whether it regulates RelA degradation by inhibiting its ubiquitination. Using a chain-specific ubiquitin antibody, we demonstrated that RelA was conjugated with lysine (K) 48-linked polyubiquitin chains in the absence of OTUB1 after αCD40 stimulation (**Fig. 3G**). These results support the idea that OTUB1 limits K48-linked ubiquitination of RelA and thereby prevents its proteasomal degradation.

In B cells, Li et al. showed that OTUB1 prevented B cell-mediated autoimmunity by stabilizing p100 and thus limiting both non-canonical and canonical NF-*Κ*B signaling. In contrast, under our experimental conditions, we did not detect increased p100 degradation in OTUB1-deficient B cells at steady state conditions or during stimulation. One possible explanation for this discrepancy is the different experimental context, including the age of the mice analyzed by Li et al.(Y. Li et al., 2019), compared with the 8–12-week-old animals used in our study. Differences in housing conditions or microbial exposure may also contribute. Rather than a dominant p100 phenotype, we detected a pronounced role of OTUB1 specifically after CD40 -mediated B cell activation. Interestingly, prolonged CD40 stimulation led to a marked loss of RelA in OTUB1-deficient B cells, while stimulation with TLR or through the BCR did not enhance a comparable degradation of RelA. These observations point to a stimulus⍰dependent role of OTUB1 in controlling which NF⍰κB subunits are engaged downstream of different receptors rather than a uniform effect on p100 stability.

Our data suggest that OTUB1 contributes to RelA stability and thereby modulates NF-κB signaling in a stimulus-dependent manner. Mechanistically, this effect is mediated at least in part through the regulation of RelA ubiquitination. OTUB1 associates with RelA, and loss of OTUB1 is accompanied by increased K48-linked ubiquitination and proteasome-dependent degradation of RelA after CD40 stimulation. Thus, RelA represents an additional OTUB1-regulated target in B cells beyond p100. Even though canonical NF-κB activation is often associated with accumulation of p100 due to the induction of *Nfkb2* gene expression, we did not observe significant differences here, supporting the idea that OTUB1 function may depend on stimulus, kinetics, or cellular contexts.

In addition to the RelA phenotype, we observed alterations in other CD40-associated upstream signaling components. OTUB1-deficient B cells displayed altered I*Κ*B*⍺* kinetics after CD40 stimulation, whereas *⍺*IgM-induced I*Κ*B*⍺* degradation was broadly comparable between genotypes. Additionally, we noted strongly reduced TRAF3 and moderately reduced TRAF6 protein levels after CD40 stimulation. Given that previous *in vitro* studies suggest that OTUB1 and OTUB2 regulate virus-induced type I interferon expression through inhibiting the ubiquitination of TRAF3 and TRAF6 within the mitochondrial RIG-I adaptor VISA (MAVS) signaling platform (S. Li et al., 2010), it is possible that OTUB1 also influences proximal CD40 signaling events in B cells, although this will require further mechanistic investigation. Together, these observations indicate that OTUB1 can shape NF⍰κB pathway output both upstream (at the level of IκB and TRAF3/6) and downstream (at the level of RelA stability), rather than simply increasing or decreasing overall NF⍰κB activation.

Functionally, in agreement with previous reports (Luo et al., 2024), OTUB1 deficiency resulted in increased marginal zone B-cell accumulation as well as enhanced proliferation in response to CD40 stimulation, and augmented class-switch responses, including increased NP-specific IgG1 production after immunization. Together, these findings identify OTUB1 as an important regulator of B-cell homeostasis and activation. Our study expands the current view of OTUB1 function in B cells by showing that, in addition to the previously described regulation of p100, OTUB1 can also constrain CD40-driven responses, at least in part by protecting RelA from K48-linked ubiquitination and proteasomal degradation. Although Luo et al. reported a reduction in both the frequency and absolute number of follicular B cells in Mb1-Cre;Otub1fl/fl mice, we observed primarily changes in follicular b cell frequencies, with total follicular B cell numbers largely preserved in Cd19-Cre;Otub1fl/fl animals, a difference that likely reflects the distinct Cre drivers and efficiencies of recombination used in the two models, because Mb1-Cre is active earlier and more uniformly in developing B cells than Cd19-Cre (Hobeika et al., 2006; Yasuda et al., 2021). Notably, this phenotype appeared to be stimulus-dependent, as we did not observe comparable alterations after LPS, BAFF, or *⍺*IgM stimulation. Thus, our findings support a context⍰specific role for OTUB1 in shaping NF⍰κB signaling and B⍰cell responses by altering the quality and temporal dynamics of NF⍰κB signaling rather than simply increasing or decreasing global NF⍰κB activation.

### Concluding Remarks

Collectively, our data show that B-cell-specific deletion of OTUB1 leads to a significant increase in marginal zone B cells and enhances CD40-driven B-cell activation and class-switching responses.

Mechanistically, we unraveled a new role for OTUB1 in the regulation of NF-κB signaling. We demonstrate that OTUB1 binds to RelA and subsequently prevents lysine-48-linked ubiquitination and degradation of this transcription factor. These results uncover a stimulus-dependent role for OTUB1 in the regulation of NF-*Κ*B signaling and establish OTUB1 as an important regulator of B-cell homeostasis.

## Data availability statement

All data generated and analyzed during this research are available.

## Ethics approval statement for human and/or animal studies

Mice were of the C57BL/6J strain, 8-12-week-old, unless otherwise stated, sex-matched male and female genders, and housed under specific pathogen-free conditions. All experiments were in accordance with the guidelines of the Translational Animal Research Center (TARC) of the University of Mainz and approved by the institutional committee on animal experimentation and the government of Rheinland-Pfalz (RLP). We have complied with all relevant ethical regulations for animal use. On the day of the experiment, animals were euthanized by gradual-fill CO₂ inhalation in accordance with institutional and national guidelines. The flow rate was set to displace approximately 30% of the chamber volume per minute. Death was confirmed by cervical dislocation.

## Clinical trial registration

NA

## Author contribution

JFV, YT, EZ, ES, and SK performed experiments, analyzed data, and prepared figures. CK prepared figures, analyzed data, and conceptualized and wrote the manuscript. DS provided the OtubFF mouse strain. NH supervised the project, wrote the manuscript, and acquired funding.

## Acknowledgments, including funding

We would like to acknowledge Petra Adams and Elena Zukowski for excellent technical help. N.H. was supported by the DFG grant HO4440/1-1, DFG grant HO4440/1-2, and Research Center Immunology (FZI) and further supported by the German Research Foundation (Project Number 318346496, SFB1292/2 TP20), and TRR355 TPA05. AW TRR355 and CRC1292.

## Supporting Information

**Supplementary Figure 1.** OTUB1 regulates B-cell differentiation and germinal center responses. **(A)** Representative histological images of spleen sections from control (Cd19-cre) and B cell-specific Otub1-deficient (Otub1BKO) mice. **(B)** Representative flow cytometry plots and quantification of T cells (CD90.2⁺) and B cells (B220⁺) in lymphoid organs of Cd19-cre and Otub1BKO mice, shown as percentage of lymphocytes and total cell numbers. **(C)** Representative flow cytometry plots and quantification of marginal zone (MZ: CD21^high^CD23^low^) and follicular (Fol: CD21^int^CD23^high^) B-cell subsets in spleens from 30-week-old Cd19-cre and Otub1BKO mice. **(D)** Serum immunoglobulin levels (IgM, IgG1, and IgG3) measured by ELISA in Cd19-cre and Otub1BKO mice. Data are shown as mean ± SEM. Each dot represents an individual mouse. Statistical significance was determined using multiple unpaired two-tailed t-tests with Welch’s correction with *p < 0.05, **p < 0.01, ***p < 0.001, ****p < 0.0001.

**Supplementary Figure 2.** Stimulus-dependent effects of OTUB1 deficiency on NF-κB and signaling pathways in B cells. **(A)** Immunoblot analysis of cytoplasmic and nuclear fractions from B cells isolated from *Cd19*-cre and Otub1BKO mice following BAFF stimulation for the indicated time points (0, 2, 4, 8, and 16h). Blots show levels of P100/P52, phospho-IκBα (p-IκBα), IκBα, RelA, and RelB. Tubulin and HDAC1 serve as cytoplasmic and nuclear loading controls, respectively. **(B)** Immunoblot analysis of cytoplasmic and nuclear extracts from B cells stimulated with LPS for the indicated time points (0, 2, 4, 8 and 16h), showing P100/P52, RelA, RelB, and c-Rel expression. Tubulin and HDAC1 serve as cytoplasmic and nuclear loading controls, respectively. **(C)** Whole-cell lysates from B cells stimulated with αCD40 for the indicated time points (0, 2, 4, 8, and 16h). Blots show phospho-IκBα (p-IκBα), IκBα, phospho-JNK (p-JNK p54/p46), total JNK, phospho-Akt at Ser473 (p-Akt (Ser473)), phospho-Akt at Thr308 (p-Akt (Thr308)), total Akt, and phospho-p38 (p-p38). Actin serves as a loading control. **(D)** Immunoblot analysis of IκBα degradation in B cells stimulated with αIgM for the indicated time points (0, 15, 30, 60, 120 min). Actin serves as a loading control. Data are representative of at least 3 independent experiments.

## Material and Methods

### Animals

Otubain^ff^ mice (Mulas et al., 2021) and the *Cd19*-cre mice (Rickert, 1997) have been described previously. Mice were analyzed at 8-12 weeks of age and were sex and age-matched. All experiments were in accordance with the guidelines of the TARC of the University of Mainz.

### Flow cytometry

Single-cell suspensions were prepared and stained with monoclonal antibodies indicated in the figure legends. Samples were acquired on FACSCanto II (Becton, Dickinson), and analyzed with FlowJo software (TriStar). Cells were treated with Fc-block (eBioscience), washed, and surface-stained.

### Cell culture and proliferation assay

8- to 12-week-old mice were used, and B cells were purified with the isolation kit (Miltenyi) using CD43 depletion or CD19^+^ MicroBeads to a purity of 95–98% as determined by flow cytometry. For the proliferation assay, 0.5x10^6^ cells were incubated with 2.5 μM cell tracer in PBS for 6 min at room temperature (RT). To stop the staining reaction, RPMI plus 10% FCS was added, and cells were washed with PBS. Final concentrations of stimuli: 10 μg/mL anti-IgM (Jackson ImmunoResearch Laboratories), 0.1 μM CpG (InvivoGen), 20 μg/mL LPS (Sigma-Aldrich) were added for 4 days. For in vitro class switch recombination, low density (0.5x 10^5^ cells/well) B cell cultures were used in DMEM medium plus 10% FCS and respective stimulation for 4 days. Final concentrations of the activating stimuli: 40 μg/mL LPS, anti-IL-4 (20 ng/mL), anti-IL-5 (1.5 ng/mL), TGF-β (2 ng/mL), BAFF (100 ng/mL), and 5 μg/mL CD40.

### Immunization and ELISA

Age-matched Otub1BKO and control mice were immunized i.p. with 10 μg NP-Ficoll (Biosearch Technologies) to measure T-cell-independent response or with 20 μg NP-CGG in alum for T-cell-dependent response. On days 7, 14, 21, and 28, serum was collected from peripheral blood. Serum Ig concentrations and NP-specific antibodies were determined by ELISA using NP-specific standards.

### Preparation of cytoplasmic and nuclear extracts for immunoblot analysis

B cells were isolated from LN and either left untreated or stimulated as indicated. Following stimulation, cells were harvested, washed twice with ice-cold PBS, and processed for subcellular fractionation. Briefly, cells were resuspended in ice-cold cytoplasmic lysis buffer containing 10 mM HEPES (pH 7.9), 10 mM KCl, 0.1 mM EDTA, 0.1 mM EGTA, and protease and phosphatase inhibitors. After incubation on ice for 15 min, NP-40 was added to a final concentration of 0.5%, and samples were vortexed briefly. Lysates were centrifuged at 12,000g for 1 min at 4°C. The supernatant containing the cytoplasmic fraction was collected and stored on ice.

The remaining nuclear pellet was washed once with cytoplasmic lysis buffer and then resuspended in nuclear extraction buffer containing 20 mM HEPES (pH 7.9), 400 mM NaCl, 1 mM EDTA, 1 mM EGTA, 1 mM DTT, and protease and phosphatase inhibitors. Samples were incubated on ice for 15–30 min with intermittent vortexing, followed by centrifugation at 12,000 g for 5 min at 4°C. The supernatant containing the nuclear fraction was collected. Protein concentrations were determined using a standard protein assay, and equal amounts of protein were subjected to SDS-PAGE and western blot analysis. For validation of fraction purity, cytoplasmic and nuclear markers were used.

### Immunoprecipitation

For immunoprecipitation, cells were lysed in IP lysis buffer (20 mM Tris pH 7.5, 150 mM NaCl, 0,5% Triton X-100, 1 mM Na_2_EDTA, 30 mM NaF, 2 mM sodium pyrophosphate, supplemented with protease inhibitor mixture (Roche Applied Science) and 1 mM N-ethylmaleimide (Sigma-Aldrich)). Lysates were normalized for protein concentration and incubated with the appropriate Abs and prepurified Dynabeads Protein G magnetic beads (Protein G was crosslinked with Bis (sulfosuccinimidyl) suberat (Thermo Fisher #21580 according to the manufacturer’s instructions) at 4°C for 16 h. After extensive washing, the bead-bound complexes were eluted using DynaMag (Invitrogen, # 12321D) in sample buffer, proteins were recovered by using Nupage LDS sample buffer (Invitrogen, #NP0007), and resolved by SDS-PAGE and immunoblotted with the appropriate antibodies. The antibodies used were as follows: anti-αβ Tubulin: cell signaling (#2148, 1:1000), anti-HDAC1: cell signaling (#34589, 1:1000), anti-Otub1: cell signaling (#3783, 1:1000), anti-K48 ubiquitin: cell signaling (#8081, 1:1000), anti-RelA: cell signaling (#8242, 1:1000), anti-p-P100: cell signaling (#4810, 1:1000), anti-p100/p52: cell signaling (#52583, 1:1000), anti-relB: Santa Cruz (#48366, 1:200)

### Transfection

Calcium phosphate transfection method was used for the transfection. 293T cells were plated in 100-mm dishes and incubated overnight at 37°C in a CO_2_ incubator. Medium was changed 6 hours prior to transfection. The CaCl_2_/DNA mix was prepared by adding 25 μg of Plasmid DNA dropwise to CaCl_2_ solution (0,2 M). After 25 min of incubation at room temperature, the CaCl_2_/DNA mix was added slowly under vortexing to 300 μl 2x Hepes-buffered saline. After 30 min of incubation at room temperature, a fine calcium phosphate/DNA precipitate was formed. The precipitates were then added, sprinkling dropwise, to the cells in the dish. After 12 hours, pre-warmed fresh medium was added. The cells were incubated at 37°C for another 16h and harvested. pRK5-HA-Ubiquitin-K48 was a gift from Ted Dawson (Addgene plasmid # 17605). Flag-HA-Outb1 was a gift from Wade Harper (Addgene plasmid # 22551). RelA cFLAG pcDNA3 was a gift from Stephen Smale (Addgene plasmid # 20012).

### Statistical analysis

For statistical analysis and graph design, Prism (GraphPad Software Inc., USA) was used. Values are presented as means ± SEM, with the number of independent experiments. Statistical differences were determined using a two-tailed Student’s *t*-test. ELISA was analyzed with SoftMax Pro 5.4.1.

## References

1. Allman, D., & Pillai, S. (2008). Peripheral B cell subsets. Current Opinion in Immunology, 20(2), 149–157. 10.1016/j.coi.2008.03.014

2. Baens, M., Fevery, S., Sagaert, X., Noels, H., Hagens, S., Broeckx, V., Billiau, A. D., De Wolf-Peeters, C., & Marynen, P. (2006). Selective Expansion of Marginal Zone B Cells in Eμ-API2-MALT1 Mice Is Linked to Enhanced IκB Kinase γ Polyubiquitination. Cancer Research, 66(10), 5270–5277. 10.1158/0008-5472.CAN-05-4590

3. Berberich, I., Shu, G. L., & Clark, E. A. (1994). Cross-linking CD40 on B cells rapidly activates nuclear factor-*kappa* B. The Journal of Immunology, 153(10), 4357–4366. 10.4049/jimmunol.153.10.4357

4. Bishop, G. A., Moore, C. R., Xie, P., Stunz, L. L., & Kraus, Z. J. (n.d.). TRAF Proteins in CD40 Signaling. In TNF Receptor Associated Factors (TRAFs) (pp. 131–151). Springer New York. 10.1007/978-0-387-70630-6_11

5. Casola, S., Otipoby, K. L., Alimzhanov, M., Humme, S., Uyttersprot, N., Kutok, J. L., Carroll, M. C., & Rajewsky, K. (2004). B cell receptor signal strength determines B cell fate. Nature Immunology, 5(3), 317–327. 10.1038/ni1036

6. Chen, Z., & Wang, J. H. (2021). How the Signaling Crosstalk of B Cell Receptor (BCR) and Co-Receptors Regulates Antibody Class Switch Recombination: A New Perspective of Checkpoints of BCR Signaling. Frontiers in Immunology, 12. 10.3389/fimmu.2021.663443

7. Chu, Y., Vahl, J. C., Kumar, D., Heger, K., Bertossi, A., Wójtowicz, E., Soberon, V., Schenten, D., Mack, B., Reutelshöfer, M., Beyaert, R., Amann, K., van Loo, G., & Schmidt-Supprian, M. (2011). B cells lacking the tumor suppressor TNFAIP3/A20 display impaired differentiation and hyperactivation and cause inflammation and autoimmunity in aged mice. Blood, 117(7), 2227–2236. 10.1182/blood-2010-09-306019

8. Coope, H. J. (2002). CD40 regulates the processing of NF-kappaB2 p100 to p52. The EMBO Journal, 21(20), 5375–5385. 10.1093/emboj/cdf542

9. Danese, S., Sans, M., & Fiocchi, C. (2004). The CD40/CD40L costimulatory pathway in inflammatory bowel disease. Gut, 53(7), 1035–1043. 10.1136/gut.2003.026278

10. Duan, B., & Morel, L. (2006). Role of B-1a cells in autoimmunity. Autoimmunity Reviews, 5(6), 403–408. 10.1016/j.autrev.2005.10.007

11. Edelmann, M. J., Iphöfer, A., Akutsu, M., Altun, M., di Gleria, K., Kramer, H. B., Fiebiger, E., Dhe-Paganon, S., & Kessler, B. M. (2009). Structural basis and specificity of human otubain 1-mediated deubiquitination. Biochemical Journal, 418(2), 379–390. 10.1042/BJ20081318

12. Franzoso, G., Carlson, L., Xing, L., Poljak, L., Shores, E. W., Brown, K. D., Leonardi, A., Tran, T., Boyce, B. F., & Siebenlist, U. (1997). Requirement for NF-κB in osteoclast and B-cell□development. Genes & Development, 11(24), 3482–3496. 10.1101/gad.11.24.3482

13. Gibb, D. R., El Shikh, M., Kang, D.-J., Rowe, W. J., El Sayed, R., Cichy, J., Yagita, H., Tew, J. G., Dempsey, P. J., Crawford, H. C., & Conrad, D. H. (2010). ADAM10 is essential for Notch2-dependent marginal zone B cell development and CD23 cleavage in vivo. Journal of Experimental Medicine, 207(3), 623–635. 10.1084/jem.20091990

14. Grossmann, M. (2000). The anti-apoptotic activities of Rel and RelA required during B-cell maturation involve the regulation of Bcl-2 expression. The EMBO Journal, 19(23), 6351–6360. 10.1093/emboj/19.23.6351

15. Hardy, R. R., & Hayakawa, K. (2015). Perspectives on fetal derived CD5 ^+^ B1 B cells. European Journal of Immunology, 45(11), 2978–2984. 10.1002/eji.201445146

16. He, M., Zhou, Z., Wu, G., Chen, Q., & Wan, Y. (2017). Emerging role of DUBs in tumor metastasis and apoptosis: Therapeutic implication. Pharmacology & Therapeutics, 177, 96–107. 10.1016/j.pharmthera.2017.03.001

17. Heltemes, L. M., & Manser, T. (2002). Level of B Cell Antigen Receptor Surface Expression Influences Both Positive and Negative Selection of B Cells During Primary Development. The Journal of Immunology, 169(3), 1283–1292. 10.4049/jimmunol.169.3.1283

18. Herhaus, L., Al-Salihi, M., Macartney, T., Weidlich, S., & Sapkota, G. P. (2013). OTUB1 enhances TGFβ signalling by inhibiting the ubiquitylation and degradation of active SMAD2/3. Nature Communications, 4(1), 2519. 10.1038/ncomms3519

19. Hobeika, E., Thiemann, S., Storch, B., Jumaa, H., Nielsen, P. J., Pelanda, R., & Reth, M. (2006). Testing gene function early in the B cell lineage in mb1-cre mice. Proceedings of the National Academy of Sciences, 103(37), 13789–13794. 10.1073/pnas.0605944103

20. Hövelmeyer, N., Reissig, S., Thi Xuan, N., Adams-Quack, P., Lukas, D., Nikolaev, A., Schlüter, D., & Waisman, A. (2011). A20 deficiency in B cells enhances B-cell proliferation and results in the development of autoantibodies. European Journal of Immunology, 41(3), 595–601. 10.1002/eji.201041313

21. Hövelmeyer, N., Schmidt-Supprian, M., & Ohnmacht, C. (2022). NF-κB in control of regulatory T cell development, identity, and function. Journal of Molecular Medicine, 100(7), 985–995. 10.1007/s00109-022-02215-1

22. Ho□velmeyer, N., Wunderlich, F. T., Massoumi, R., Jakobsen, C. G., Song, J., Wo□rns, M. A., Merkwirth, C., Kovalenko, A., Aumailley, M., Strand, D., Bru□ning, J. C., Galle, P. R., Wallach, D., Fa□ssler, R., & Waisman, A. (2007). Regulation of B cell homeostasis and activation by the tumor suppressor gene *CYLD*. The Journal of Experimental Medicine, 204(11), 2615–2627. 10.1084/jem.20070318

23. Komander, D., Clague, M. J., & Urbé, S. (2009). Breaking the chains: structure and function of the deubiquitinases. Nature Reviews Molecular Cell Biology, 10(8), 550–563. 10.1038/nrm2731

24. Li, S., Zheng, H., Mao, A.-P., Zhong, B., Li, Y., Liu, Y., Gao, Y., Ran, Y., Tien, P., & Shu, H.-B. (2010). Regulation of Virus-triggered Signaling by OTUB1- and OTUB2-mediated Deubiquitination of TRAF3 and TRAF6. Journal of Biological Chemistry, 285(7), 4291–4297. 10.1074/jbc.M109.074971

25. Li, Y., Yang, J.-Y., Xie, X., Jie, Z., Zhang, L., Shi, J., Lin, D., Gu, M., Zhou, X., Li, H. S., Watowich, S. S., Jain, A., Yun Jung, S., Qin, J., Cheng, X., & Sun, S.-C. (2019). Preventing abnormal NF-κB activation and autoimmunity by Otub1-mediated p100 stabilization. Cell Research, 29(6), 474–485. 10.1038/s41422-019-0174-3

26. Luo, V. M., Shen, C., Worme, S., Bhagrath, A., Simo-Cheyou, E., Findlay, S., Hébert, S., Wai Lam Poon, W., Aryanpour, Z., Zhang, T., Zahedi, R. P., Boulais, J., Buchwald, Z. S., Borchers, C. H., Côté, J.-F., Kleinman, C. L., Mandl, J. N., & Orthwein, A. (2024). The Deubiquitylase Otub1 Regulates the Chemotactic Response of Splenic B Cells by Modulating the Stability of the γ-Subunit Gng2. Molecular and Cellular Biology, 44(1), 1–16. 10.1080/10985549.2023.2290434

27. Luong, L. A., Fragiadaki, M., Smith, J., Boyle, J., Lutz, J., Dean, J. L. E., Harten, S., Ashcroft, M., Walmsley, S. R., Haskard, D. O., Maxwell, P. H., Walczak, H., Pusey, C., & Evans, P. C. (2013). Cezanne Regulates Inflammatory Responses to Hypoxia in Endothelial Cells by Targeting TRAF6 for Deubiquitination. Circulation Research, 112(12), 1583–1591. 10.1161/CIRCRESAHA.111.300119

28. Maillard, I., Weng, A. P., Carpenter, A. C., Rodriguez, C. G., Sai, H., Xu, L., Allman, D., Aster, J. C., & Pear, W. S. (2004). Mastermind critically regulates Notch-mediated lymphoid cell fate decisions. Blood, 104(6), 1696–1702. 10.1182/blood-2004-02-0514

29. Mulas, F., Wang, X., Song, S., Nishanth, G., Yi, W., Brunn, A., Larsen, P.-K., Isermann, B., Kalinke, U., Barragan, A., Naumann, M., Deckert, M., & Schlüter, D. (2021). The deubiquitinase OTUB1 augments NF-κB-dependent immune responses in dendritic cells in infection and inflammation by stabilizing UBC13. Cellular & Molecular Immunology, 18(6), 1512–1527. 10.1038/s41423-020-0362-6

30. Tavares, R. M., Turer, E. E., Liu, C. L., Advincula, R., Scapini, P., Rhee, L., Barrera, J., Lowell, C. A., Utz, P. J., Malynn, B. A., & Ma, A. (2010). The Ubiquitin Modifying Enzyme A20 Restricts B Cell Survival and Prevents Autoimmunity. Immunity, 33(2), 181–191. 10.1016/j.immuni.2010.07.017

31. Xue, L., Morris, S. W., Orihuela, C., Tuomanen, E., Cui, X., Wen, R., & Wang, D. (2003). Defective development and function of Bcl10-deficient follicular, marginal zone and B1 B cells. Nature Immunology, 4(9), 857–865. 10.1038/ni963

32. Yasuda, T., Saito, Y., Ono, C., Kawata, K., Baba, A., & Baba, Y. (2021). Generation and characterization of CD19-iCre mice as a tool for efficient and specific conditional gene targeting in B cells. Scientific Reports, 11(1), 5524. 10.1038/s41598-021-84786-6

33. Zhou, H., Liu, Y., Zhu, R., Ding, F., Cao, X., Lin, D., & Liu, Z. (2018). OTUB1 promotes esophageal squamous cell carcinoma metastasis through modulating Snail stability. Oncogene, 37(25), 3356–3368. 10.1038/s41388-018-0224-1

