## Supplementary figures and images for "OTUB1 controls marginal zone B-cell development by stabilizing RelA in a CD40-dependent manner"

### Supplementary Figure 1

#### Supplementary Figure 1: OTUB1 regulates B cell differentiation

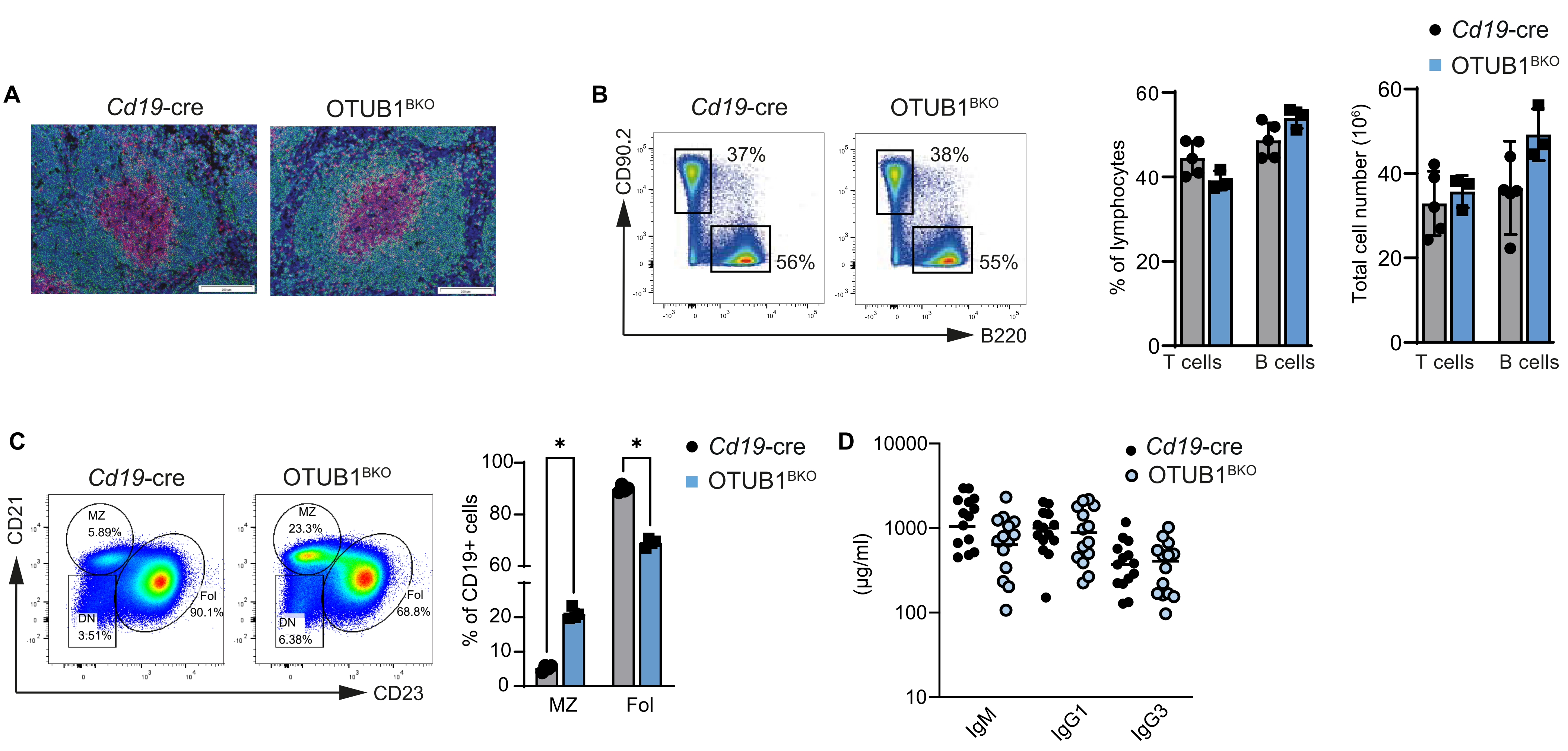
