## Supplementary Figure 2 for "OTUB1 controls marginal zone B-cell development by stabilizing RelA in a CD40-dependent manner"

Supplementary Figure 2: Stimulus-dependent effects of OTUB1 deficiency on NF-κB and signaling pathways in B cells

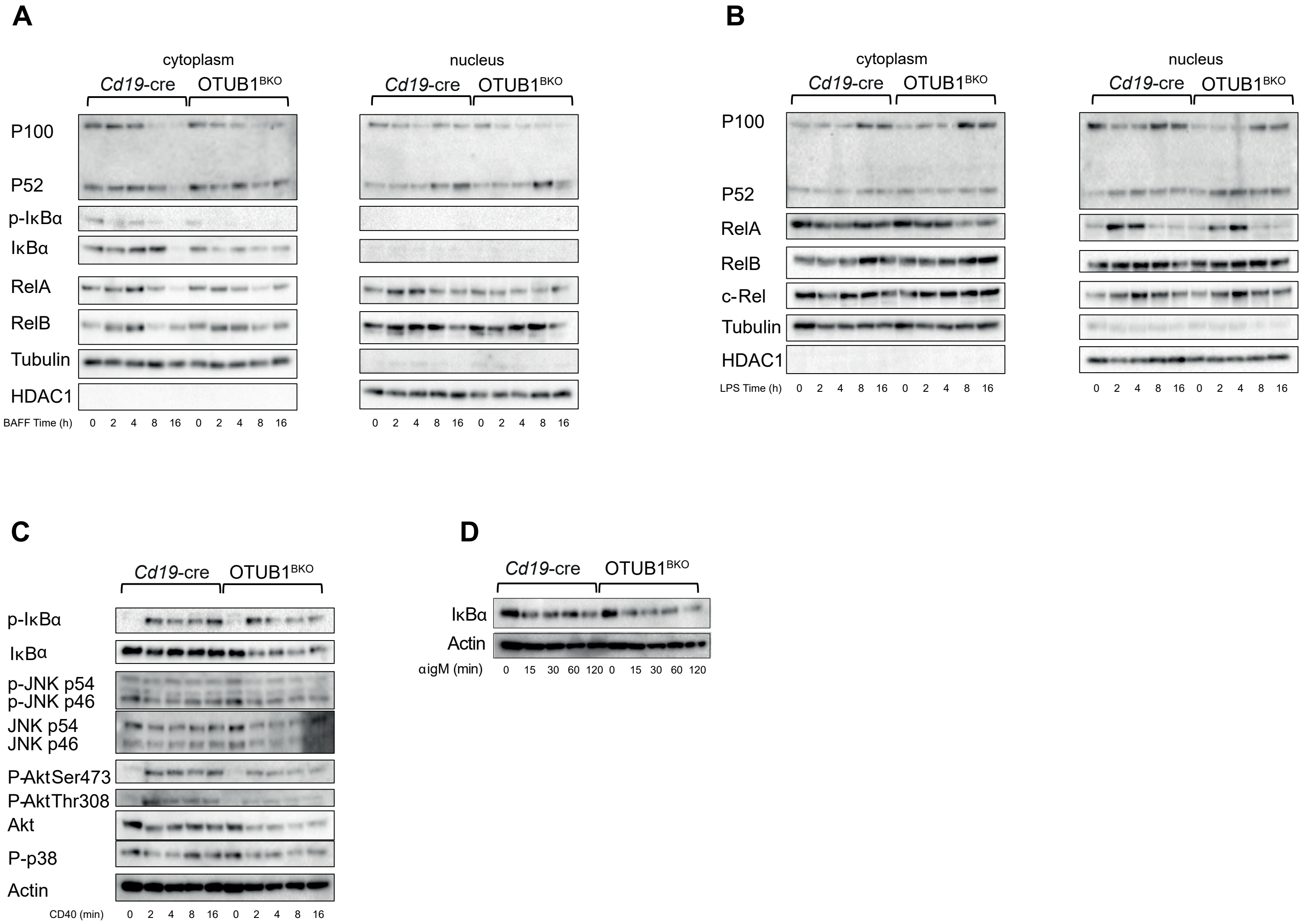
